# A non-invasive radar system for automated behavioural tracking: application to sheep

**DOI:** 10.1101/2020.12.09.418038

**Authors:** Alexandre Dore, Cristian Pasquaretta, Dominique henry, Edmond Ricard, Jean-François Bompard, Mathieu Bonneau, Alain Boissy, Dominique Hazard, Hervé Aubert, Mathieu Lihoreau

## Abstract

Automated quantification of the behaviour of freely moving animals is increasingly needed in ethology, ecology, genetics and evolution. State-of-the-art approaches often require tags to identify animals, high computational power for data collection and processing, and are sensitive to environmental conditions, which limits their large-scale utilisation. Here we introduce a new automated tracking system based on millimetre-wave radars for real time robust and high precision monitoring of untagged animals. To validate our system, we tracked 64 sheep in a standard indoor behavioural test used for genetic selection. First, we show that the proposed radar application is faster and more accurate than conventional video and infrared tracking systems. Next, we illustrate how new behavioural estimators can be derived from the radar data to assess personality traits in sheep for behavioural phenotyping. Finally, we demonstrate that radars can be used for movement tracking at larger spatial scales, in the field, by adjusting operating frequency and radiated electromagnetic power. Millimetre-wave radars thus hold considerable promises for high-throughput recording of the behaviour of animals with various sizes and locomotor modes, in different types of environments.

## 1. Introduction

Animal behaviour research increasingly requires automated recording and analyses of movements (Branson et al., 2009). The emerging field of computational ethology provides methods for high-throughput monitoring and statistical analyses of movements that enable the quantitative characterisation of behaviour on large numbers of individuals, the discovery of new behaviours, but also the objective comparison of behavioural data across studies and species (Anderson and Perona, 2014; Brown and de Bivort, 2018).

These quantitative approaches are particularly powerful to study inter-individual behavioural variability or personalities in animal populations (Morand-Ferron et al., 2015). In livestock, for instance, large-scale genetic selection programmes are based on the measurements of several hundreds (if not thousands) of farm animals (O’Brien et al. 2014). Many behavioural tests have been developed to assess behavioural and personality traits in farm animals (Canario et al. 2013), and some applications have been developed in breeding programmes for instance to discard the more aggressive individuals for beef cattle production (Phocas et al. 2006). Behavioural measures are frequently obtained from direct observations by the experimenters or farmers (e.g., Boissy et al., 2005), limiting use of behavioural criteria for breeding programmes. Indeed, the absence of automated measurements make data collection cumbersome, time-consuming and prone to biases, which currently limits the ability to quantify behavioural traits at the experimental or commercial farm level.

Tracking methods involving on-board devices, such as Global Positioning Systems (GPS) (Tomkiewicz et al., 2010), radio telemetry (Cadahia et al., 2010), radio frequency identification (RFID) (Voulodimos et al., 2010) or harmonic radar (Riley et al., 1996), are hardly suitable for detailed high throughput behavioural phenotyping due to the limited accuracy and duration of measurements. Best available methods therefore involve image-based analyses (Pérez-Escudero et al., 2014; Romero-Ferrero et al., 2019). However, these techniques often require large computational resources to acquire and process images (Garcia et al 2019) and are sensitive to light variation (Dell et al., 2014).

Recently, Frequency-Modulated Continuous-Wave (FMCW) radars operating in the millimetre-wave frequency band have been proposed for automated tracking of animal behaviour (sow: Dore et al., 2020b, bees: 2020a; sheep: Henry et al., 2018). In this approach, it is possible to record the 1D movements (distance to radar) of individual sheep in an arena test (Henry et al., 2018). Tracking with FMCW radars has the great advantage of being non-invasive (does not require a tag), insensitive to light intensity variations, and fast (as it does not require large memory resource). FMCW radars therefore provide considerable advantages for the development of automated high-throughput analyses in regard to more conventional approaches (e.g. video and infrared tracking systems).

Here we developed a millimetre-wave FMCW radar system for automated tracking and analyses of the 2D trajectories of freely moving animals. We illustrated our approach by analysing the behaviour of 64 sheep in an “arena test” commonly used to estimate the sociability of individual sheep in genetic selection (Boissy et al., 2005; Hazard et al., 2014). First, we compared the speed and accuracy of radar tracking with more conventional video and infrared tracking systems. We then derived new behavioural estimators computed from the radar data, that could be used for large-scale behavioural phenotyping. Finally, we tested the radar system for long-distance tracking, in the field, by adjusting radar emission frequency and radiated electromagnetic power.

## 2. Materials and methods

### Sheep

We ran the experiments in July 2019 at the experimental farm La Fage of the French National Research Institute for Agriculture, Food, and Environment (INRAE), France (43.918304, 3.094309). We tested 64 lambs (32 males, 32 females) *Ovis aries* with known weight (range: 12kg – 31.3kg) and age (range: 59 days – 88 days). Ewes and their lambs were reared outdoor on rangelands. After weaning, lambs were reared together outside and tested for behaviour 10 days later. This delay allowed the development of social preferences for conspecifics instead of preference for mother. All the lambs were previously tested in a “corridor test” to estimate their docility towards a human. Briefly the test pen consisted of a closed, wide rectangular circuit (4.5 × 7.5 m) with opaque walls (Boissy et al., 2005). A non-familiar human entered the testing pen and moves at constant speed through the corridor until two complete tours had been achieved. Every 5 s (i.e. the corridor was divided into 6 virtual zones and 1 zone was crossed every 5 s by the human) the zones in which the human and the animal were located were recorded and the mean distance separating the human and the lamb was calculated. The walking human also recorded with a stopwatch the total duration when the human saw the head of the lamb to discriminate between fleeing and following lambs. The reactivity criteria towards an approaching human was constructed by combining both measurements (for more details see Hazard et al., 2016). The higher was the resulting variable (i.e. docility variable), the more docile was the animal.

### Arena test

We tested all the sheep in the arena test, a standard protocol for assessing the sociability of sheep through measures of inter-individual variability in social motivation in the absence or presence of a shepherd (Boissy et al., 2005; Hazard et al., 2014). Briefly, a sheep (focal sheep) was introduced in the pen (2mx7m) with artificial lights (Fig. 1A; for more details see Ligout et al. 2011). Three other sheep from the same cohort (social stimuli) were placed behind a grid barrier, on the opposite side of the corridor entrance. The test involved three phases (Fig. 1B). In phase 1, the focal sheep could explore the corridor for 15s and see its conspecifics through a grid barrier. In phase 2, visual contact between the focal sheep and the social stimuli was disrupted using an opaque panel pulled down from the outside of the pen. This phase was used to assess the sociability of the sheep towards its conspecifics and lasted 60s. In phase 3, visual contact between the focal sheep and its conspecifics was re-established and a man was standing still in front of grid barrier for 60s. This phase was used to assess the sociability of the focal sheep towards conspecifics in presence of a motionless human.

**Figure 1:**
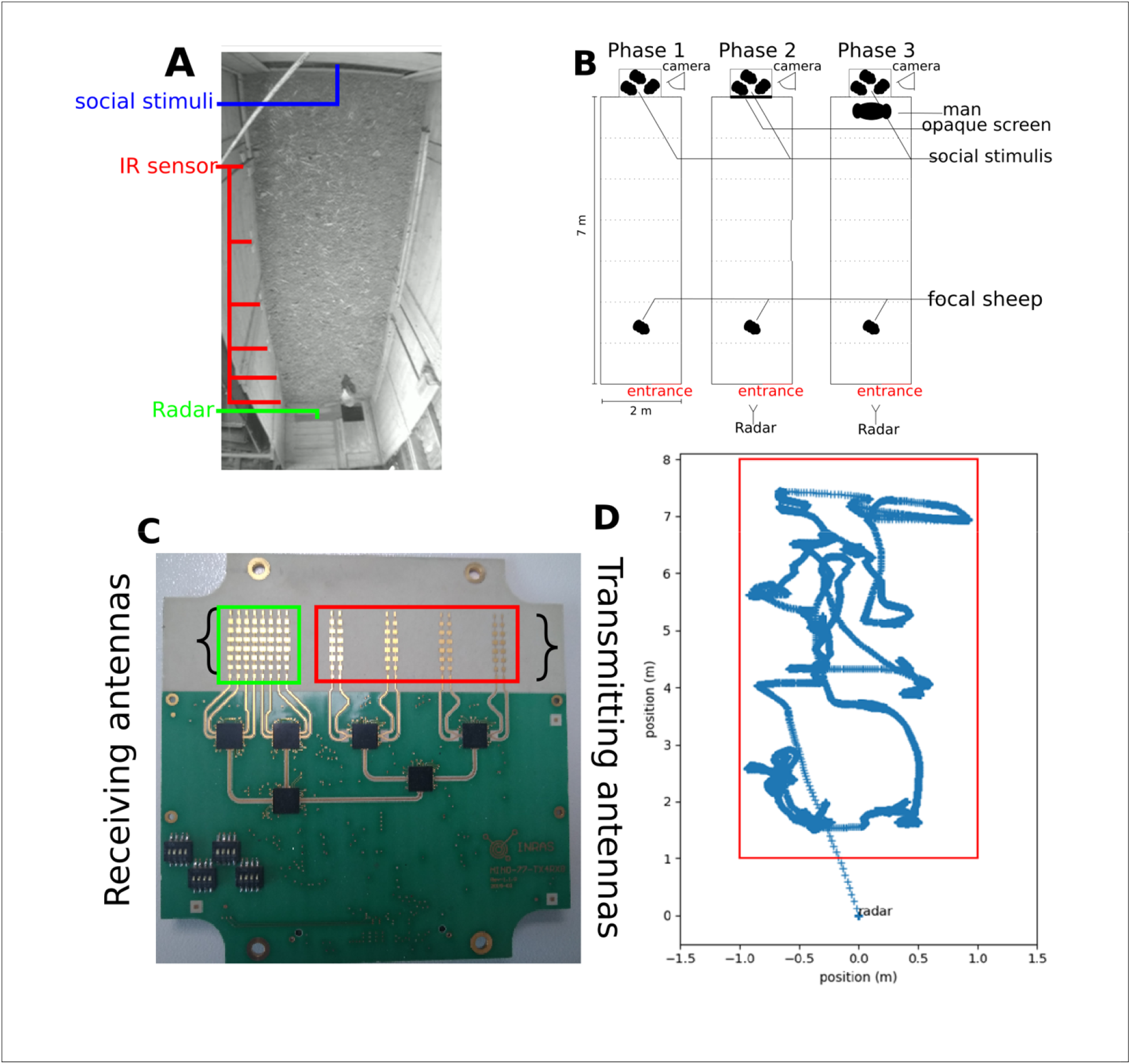
Corridor test. A. Top view of the focal sheep and the social stimuli in the corridor (example image extracted from video data). B. schematic representation of experiments phases 1, 2 and 3. C. Image of the FMCW radar frontend (phot credit AD). Each rectangle corresponds to rectangular patch (Haderer et al., 2008). D. Example of position estimations of a sheep over time after removing the clutter and normalizing the estimated value.

### Data collection

We measured the displacement of the focal sheep in phases 2 and 3 of the arena test (phase 1 is the initiation phase) using three automated tracking systems: infrared sensors, a video camera, and a millimetre-wave FMCW radar. During the measures, an experimenter also recorded the number of high-pitched bleats by the focal sheep, a proxy of the sociability (i.e. sociability variable) of the sheep (Boissy et al., 2005).

#### Tracking with infrared cells

Infrared cells were previously used to quantify sheep behaviour in the arena test. We placed two infrared sensors every meter along the arena length (Fig. 1A). We used two infrared sensors to determine the direction of the moving sheep. The focal sheep was recorded each time it passed through one of these sensors. Sheep movements were thus tracked in 1D (longitudinal movements in the corridor) and data resolution was 1m.

#### Video tracking

We placed a video camera on one end of the arena (opposite to entrance side, Fig. 1B). The camera was elevated 2m above ground in order to film the entire arena. Sheep movements were tracked in 2D. For image processing, we applied a detection algorithm using the state-of-the-art image object detector tiny-YOLO (You Only Look Once) network, which is a version of the YOLO model adapted for faster processing allowing 244 images of 0.17 mega pixels (416 × 416 pixels) per second (on a TITAN × graphics card) (Redmon et al., 2016). This neural network was pre-trained on the PASCAL Visual Object Classes Challenge dataset (Everingham et al., 2012).

#### Radar tracking

We placed a millimetre-wave FMCW radar (Fig. 1C, see technical characteristics in Table 1) at one end of the arena (entrance side, Fig. 1B). The radar was setup outside of the test pen behind a Styrofoam wall transparent to millimetre-wave (Dietlein et al., 2008). The transmitting antenna array radiated a repetition over time of a so-called chirp (i.e. a saw-tooth frequency-modulated signal (Balanis, 2011)). The chirp was backscattered by the targeted focal sheep, but also by the surrounding scene which provides undesirable radar echoes called the electromagnetic clutter. The total backscattered signal was then collected by the receiving antenna array and processed to mitigate the clutter and to derive the sheep 2D trajectory from radar data. In the millimetre-wave frequency range, the detectability of the sheep depends mainly on the bandwidth of the frequency modulation, the beamwidth of the radar antennas, and the radiated electromagnetic power (Balanis, 2011)).

**Table 1:** Technical characteristics of the FMCW radar used for indoor tracking (Haderer et al., 2008) and outdoor tracking (Simon et al., 2014).

| Name | Indoor tracking | Outdoor tracking | Note |
| --- | --- | --- | --- |
| Operating frequency | 77GHz | 24GHz | This frequency is also called the *carrier frequency* of the frequency-modulated signal transmitted by the radar |
| Modulation Bandwidth | 3GHz | 800MHz | Frequency interval, centred at the operating frequency, used for the saw-tooth frequency modulation of the transmitted signal |
| Resolution | 5cm | 18.75cm | Resolution = $\frac{c}{2B}$, with c the celerity and B the modulation bandwidth |
| Ramp time | 256 μsec | 1ms | Up-ramp duration of the saw-tooth frequency-modulated signal (or chirp duration) |
| Repetition time | 50ms | 30ms | Period of the transmitted frequency-modulated signal (or chirp repetition interval) |
| Number of linear arrays in the transmitting array antenna | 4 | 1 | One linear array composed of 8x2 rectangular patches radiating elements |
| Number of linear arrays in the receiving array antenna | 8 | 2 | Eight linear arrays composed of 8 rectangular patches radiating elements |
| Main lobe beamwidth of the transmitting array antenna in the horizontal plane | 50° | 58° | Angular range (or field of view) of the radar illumination in the horizontal plane |
| Transmitted power | 100mW | 100mW | Power delivered at the input terminals of the transmitting array antenna (the radiated power is defined as the product of the transmitted power by the efficiency of the antenna) |

### Radar signal processing

Processing of radar data included two main steps. First, we extracted the position of the animal. Next, we computed behavioural parameters to characterize the movement of the animal.

#### Extraction of sheep positions

We extracted the distance of the focal sheep to the radar and its direction in the horizontal plane of the scene. To mitigate the electromagnetic clutter, we estimated the mean value (*mean*) and standard deviation (*std*) of the signal provided by the radar in absence of the sheep and derived the signal *Detection*(*r, θ*) from the signal S_*radar*_(*r, θ*) delivered in presence of the animal, as follows:

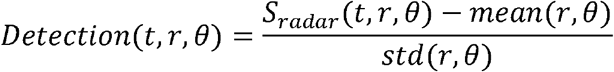

Where *r* is the distance to the sheep, *mean* is the time-averaged signal radar coming from the range *r* and angular position *θ*, *std* is the time-standard deviation of the radar signal. Fig. 1D shows an example of position estimations of a sheep over time after removing the clutter.

#### Extraction of new behavioural parameters

From the 2D trajectory data, we extracted new behavioural parameters to characterize sheep movements using three approaches.

##### Computation of behavioural classes

We statistically identified broad classes of behaviour using automated classification. First, we extracted movement parameters (speed, sinuosity and speed of displacement) from the trajectories. Then, to differentiate the movement modifying the distance of the sheep from the social stimuli (i.e. the three conspecifics) and a lateral movement, the speed vector was split in two dimensions: along the two lateral walls of the corridor and across the two longitudinal walls. These characteristics were estimated on time windows of 1s for each sheep and for each experimental phase. Finally, to derive behavioural classes, we performed an extraction from a Gaussian mixture model using the extracted data for each lamb (Reynolds and Rose, n.d.). The number of classes (i.e. the number of Gaussians to be used) was determined by comparing models using 1 to 15 classes. We selected the model with the lowest Akaike score which represents the model with the features best explaining the parameter under consideration (Burnham and Anderson, 2002).

##### Computation of behavioural transitions

We determined behavioural changes over time using Ricker wavelet processing (Ryan, 1994; see examples Fig. 3). Wavelet processing consists in filtering the sheep position signal using a wavelet as a filter (Poirier et al., 2009). This type of filtering is applied to several time scales, thus allowing the detection of a behaviour regardless of how long this behaviour lasts. Our aim was to determine the precise moments when the focal sheep changed its moving behaviour, which was estimated using the spectrum described by each wavelet. We observed that the number of local maxima in the wavelet transform coefficients is sensitive to the number of behaviour changes (see example in Fig. S1B).

**Figure 3:**
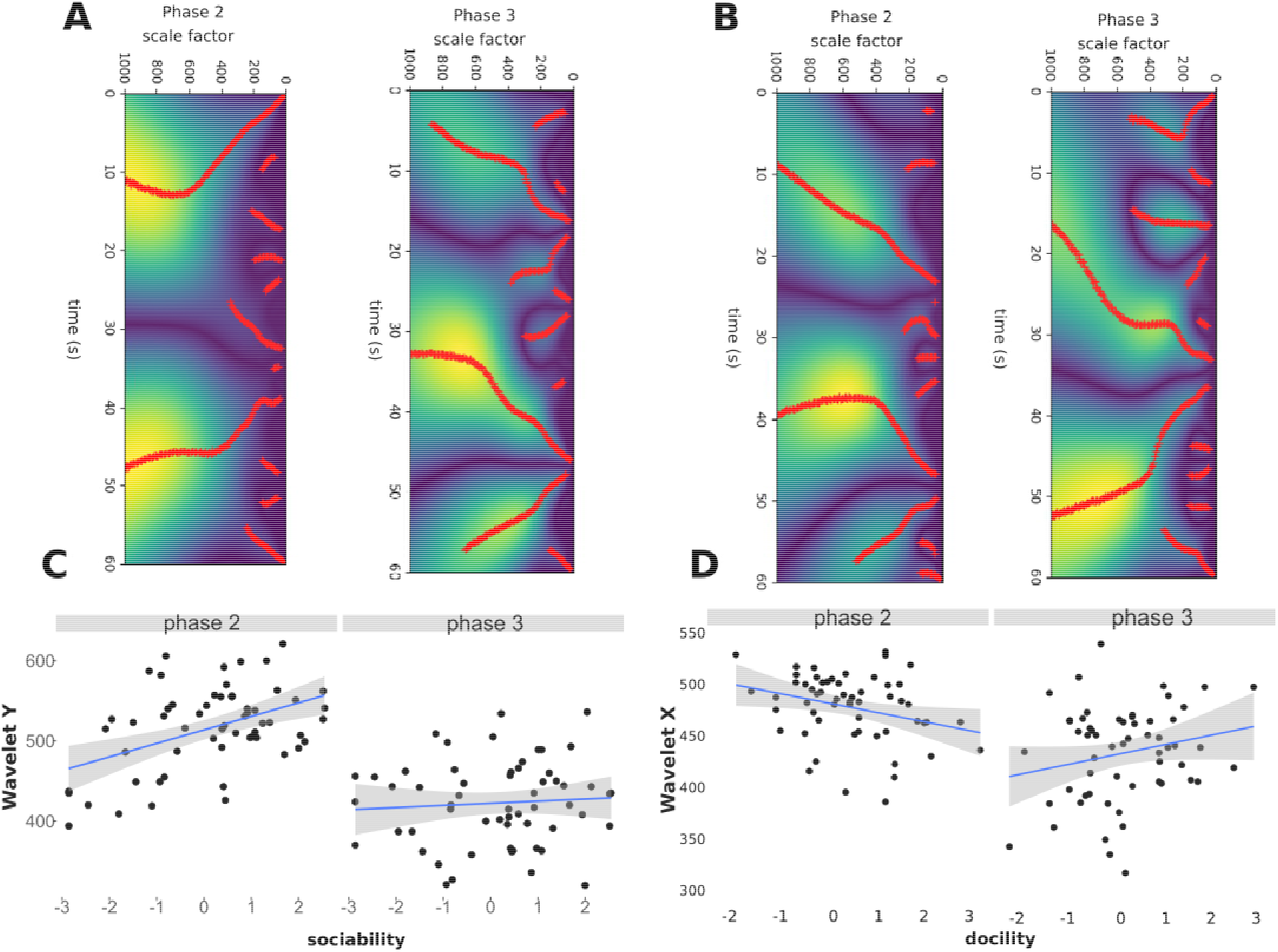
Wavelet analyses. A. Example of wavelet transform for lateral movements (X). Red dots correspond to the detection of a change in the displacement at scale factor and time position (i.e., a local maximum of the wavelet transform of the signal position). B. Example of wavelet transform for longitudinal movements (Y). C. Relationship between the number of local maxima (red dots in Figures A and B) in the wavelet extraction and the degree of sociability of sheep during phases 2 and 3. D. Relationship between the number of wavelets and the degree of docility of sheep during phases 2 and 3. See details of models in Table 4. N = 64 sheep.

##### Computation of space coverage

We investigated the space occupied across time by the focal sheep using heatmaps (see examples Fig. 4). We partitioned the arena into 80 zones of 44×40cm each (i.e. 16 partitions along the arena length and 5 partitions along the arena width). We chose this zone dimension because it corresponded to the width of a small lamb (Idris et al., 2011). We counted the number of zones the focal sheep remained in for more than 200ms. We considered that a lamb stayed in a given zone for less than 200ms either because it was positioned at the edge of this zone or because the estimation of position by the radar was inaccurate (this situation occurred for less than 10% of detections).

**Figure 4:**
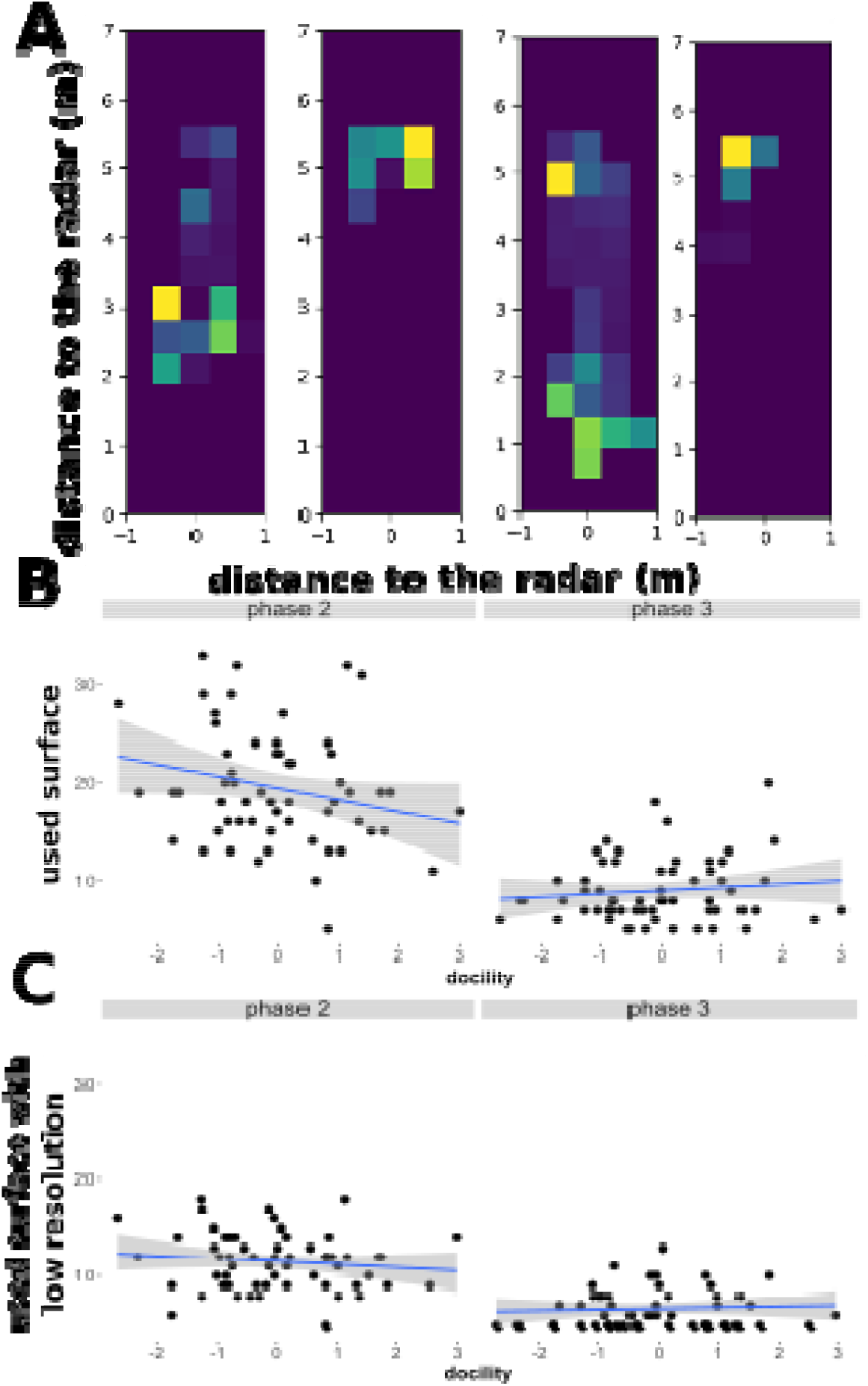
Heatmap analyses. Relationship between the number of zones occupied by the lambs and the degree of docility in phase 2 and phase 3. A. Low spatial resolution grid (cell dimension: 0.6 × 1m). B. High spatial resolution grid (cell dimension: 0.44 × 0.40 m). Top: examples of heatmaps. Bottom: Correlations. See details of models in Table 6. N = 64 sheep.

### Outdoor radar tracking

We ran outdoor experiments in order to demonstrate the applicability of our radar system for long-range tracking of sheep (Fig. 4). These measurements where done in an open space with no obstacles (60m × 15m). A man moved in order to induce animal movements. We tested one female sheep (2 years, 60 Kg). To enhance detection range to 40 m, we used a FMCW radar with a lower operating frequency (24GHz; (Simon et al., 2014)). At constant transmitted power, lower frequencies enables to reduce the free-space attenuation of the radiated electromagnetic power (Balanis, 2011). We used the same signal processing as with the 77GHz radar.

### Statistical analyses

We ran all analyses using the programming environment R(R Core Team, 2014). Raw trajectory data extracted from radar and video measures are available in Dataset S1.

#### Comparison of the performances of the different tracking systems

We tested the ability of radar and video tracking systems to capture the same information as the infrared cells from the computation of two parameters: (1) a crossing rate and (2) a proximity score (respectively LOCOM and PROX in Hazard et al. 2014). The crossing rate is the number of times the animal moved between virtual zones defined by the infrared cells (1×2m) without distinguishing between the zones nor the direction of movement. This rate thus provides information on the displacement activity of animals seeking contact with their conspecifics. The proximity score is the total duration the focal sheep remained in each zone weighted as the animal moves closer to its conspecifics. The weight depends on the distance between the sheep and the conspecifics:

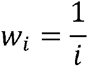

with *w_i_* the weight of the zone considered, *i* the index of the zone delimited by the infrared sensors. *i* = 1 corresponds to the zone close to the conspecifics, and *i* = 7 corresponds to the zone close to the corridor entrance. Thus, an animal with a high proximity score spent more time close to its conspecifics. We calculated these two parameters from data collected from infrared cells, video and radar tracking systems and compared them using Pearson’s correlation tests (using ‘stats’ R package).

We compared the accuracy of radar and video data systems by running a general linear mixed model (GLMM using glmer function in ‘lme4’ R package (Bates et al., 2014)) testing the effect of the tracking method on the proportion of false detections, with sheep identity as random factor. We estimated the correlation between the number of false detections with the two methods using the Pearson’s correlation test.

#### Analyses of new movement features

We tested the influence of the sheep characteristics on behavioural classes using generalised models (GLMMs). Models (binomial family) tested the effects of phase, docility, weight, age, sociability, sex, and dual interactions of each variable with test phase, on the proportion of time spent in fast movements (behavioural class 2). Interactions between more than two variables were excluded because of the high number of variables relative to sample size. Sheep identity was included as a random factor. We ran a model selection by using all features combinations (age, weight, docility, sociability, sex and the phase when the radar measurement was done). We kept the model with the highest Akaike score. We used a similar procedure to test the influence of the sheep characteristics of the lambs on continuous wavelet transforms (Gaussian models) and heatmaps (Poisson models). For the continuous wavelet transform, we performed two different wavelet transforms on the lateral and on the longitudinal position of the sheep. For heatmaps, we tested different grid resolutions characterised by zone sizes: a low resolution grid with 21 zones (3 × 7) and a high resolution grid with 45 zones (5 × 15).

#### Classification of behavioural types

To classify sheep from on our new behavioural estimators, we ran a principal component analysis (PCA) based on the eight behavioural measures extracted from the radar data in phase 2 and phase 3: proportion of fast movements (class 1) out of all movements (class 1 + class 2), longitudinal movements (wavelets Y), latitudinal movements (wavelets X), space coverage (heatmaps).

## 3. Results

### Radar tracking is more faster and more accurate than video tracking

To validate the radar tracking system, we compared the data obtained from the infrared cells, video and radar. We analysed data from 58 (29 males, 29 females) out of the 64 sheep initially tested, because some recordings failed during measurements due to human errors or miss detection by infrared cells.

Both data collected by radar and video enabled to capture information given by infrared cells with high fidelity. Proximity scores and crossing rates obtained from infrared cells were positively correlated with data obtained from radar (Pearson correlation; proximity: r = 0.77, p < 0.001; crossing rate: r = 0.87, p < 0.001) and video (Pearson correlation test; proximity: r = 0.91, p < 0.001; crossing rate: r = 0.34, p = <0.001). Imperfect correlation between proximity scores and crossing rates obtained from different tracking methods are caused by the different reference points used for tracking: infrared cells detect the full body of the sheep, whereas image analysis detects the centre of mass of the sheep, and the radar detects body parts of the sheep that are closest to it.

The comparison between the radar and the video data showed both tracking systems involved low levels of false detections (i.e. when the distance between the body centre of the sheep and its estimated position is greater than one sheep body length). However, radar tracking generated ca. three times less false detections than video tracking (Binomial GLMM, z = −3.595, p < 0.001; Table 2). False detection with the two methods had different origins. False video detections resulted from insufficient colour contrast between the lamb and the background (e.g. when the lamb was close to the wall of the corridor), whereas false radar detections resulted from the low angular resolution of the radar and potential multiple reflections of the transmitted electromagnetic signal by obstacles in the scene (such as walls, ground, human). Radar and video tracking systems are therefore complementary. The combination of the two systems gave the exact position of the sheep in 97.83% of the measures.

**Table 2:** Comparison of data processing characteristics with radar and video tracking systems.

| Tracking method | Radar | Video |
| --- | --- | --- |
| Number of measures per second | 50 | 25 |
| Read Only Memory (ROM) for all measures of a sheep | 151Mo | 62Mo |
| Random Access Memory (RAM) per measure | 524Kb | 3.7Mb |
| Processing time per measure | <20ms | 250ms |
| Total false detection rate | 6% | 16% |

Radar tracking had additional advantages over video tracking in terms of data processing (Table 2). The radar produced two times more measures per second. Radar processing was also much faster and therefore it may be used for real time data analyses. Radar measures were of similar size as video measures (ROM), but required ca. 7 times less memory (RAM) to process. Finally, radar processing did not require a learning phase with important data collection and a time-consuming training phase that can last several hours just for the adaptation of the model, or several days if the network is not trained beforehand.

### New behavioural indicators from radar data

The following analyses were made on the 64 sheep tested (32 males, 32 females). The obtained 2D trajectory data offered the opportunity for high resolution analyses of sheep movements.

#### Behavioural classes: detection of slow and fast movements

We applied a Gaussian Mixture Model (GMM) procedure to statistically identify behavioural classes from the 2D trajectory data. We found four behavioural classes (Fig. 2A). Class 1 (51.3% of measures) was characterized by null or slow movements (“slow movements”). Class 2 (35.48% of measures) was characterized by fast movements with low sinuosity (“fast movement”). Class 3 (10.2% of measures) was characterized by fast movements with high sinuosity (“fast tortuous”). Class 4 (3.01% of measures) was characterized by slow movements with high sinuosity (“slow tortuous”). Each of the two behavioural classes with strong sinuosity (classes 3 and 4) represented less than 10% of all data. We thus focused our analyses on slow and fast movements (classes 1 and 2).

**Figure 2:**
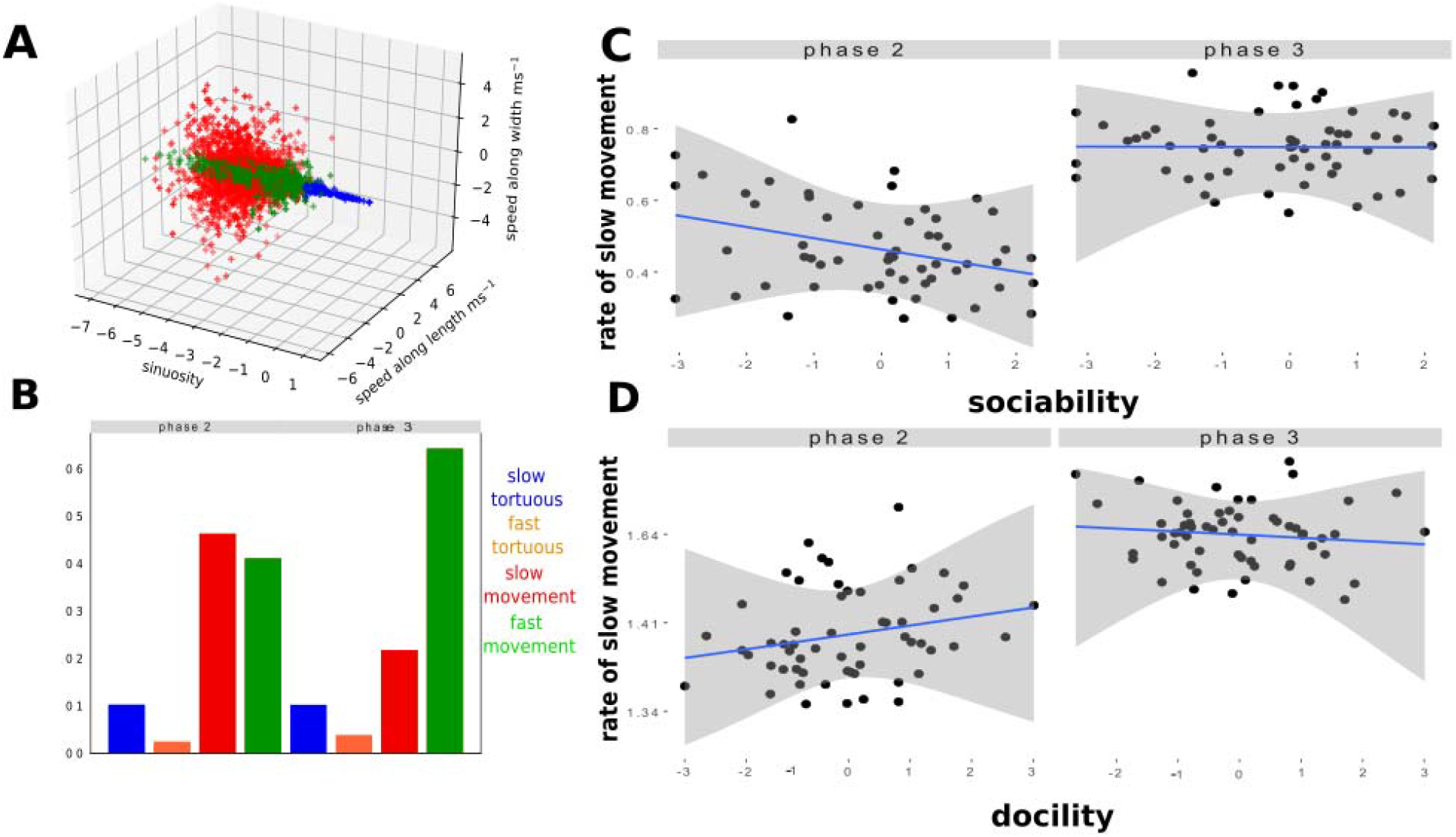
Analyses of behavioural classes. A. Distribution of the four behavioural classes after a Gaussian Mixture Model. B. Frequency of behavioural classes during phase 2 and phase 3 of the corridor test. C. Correlation between the proportion of time spent in slow movements and the sociability score of sheep during phase 2 and 3 (see details of models in Table 3). D. Correlation between the proportion of time spent in slow movements and the sociability score of sheep during phase 2 and 3 (Table 3). N = 64 sheep.

We tested the effects of the individual characteristics of sheep on time spent in each behavioural class using GLMMs. Results of the best model (i.e. highest Akaike score) are summarized in Table 3 (see model selection in Table S1). Male, old, large and highly docile sheep spent significantly more time in slow movement during phase 3 than during phase 2 (Fig. 2C). Highly sociable sheep also spent more time in slow movements during phase 3 than during phase 2 (Fig. 2D).

**Table 3:** Analyses of behavioural classes. Results of the best GLMM (binomial family, after model selection – see Table S1). The model tested the effects of phase, docility, weight, sex, age, sociability, and dual interaction of each variable with phase, on the proportion of time spent in fast movements (behavioural class 2). Sheep identity was included as a random factor. Significant effects (p<0.05) are shown in bold.

| Variable | Estimate | z-value | P-value |
| --- | --- | --- | --- |
| **Phase** | **-1.272453** | **-102.321** | **< 0.001** |
| **Docility** | **-0.134811** | **-2.725** | **0.006** |
| Weight | 0.042312 | 0.753 | 0.451 |
| Sex | -0.018224 | -0.165 | 0.869 |
| Age | -0.095193 | -1.673 | 0.094 |
| **Sociability** | **0.137659** | **3.596** | **< 0.001** |
| **Phase x docility** | **0.156352** | **19.868** | **< 0.001** |
| **Phase x weight** | **0.046431** | **5.223** | **< 0.001** |
| **Phase x sex** | **0.068090** | **3.882** | **< 0.001** |
| **Phase x age** | **0.078200** | **8.537** | **< 0.001** |
| **Phase x sociability** | **-0.127900** | **-20.900** | **< 0.001** |

#### Wavelet analyses: detection of erratic behavioural transitions

We quantified behavioural changes during time (variation in speed, direction, or both) using continuous wavelet analyses. We tested the effects of the individual characteristics of sheep on the frequency of behavioural changes using GLMMs and model selection (Tables S2 and S3). When considering longitudinal movements along the corridor length (Table 4A), we found that males made more behavioural transitions in phase 3 than in phase 2. Highly sociable sheep made more behavioural transitions in phase 3 than in phase 2 (Fig. 3A). When considering latitudinal movements along the corridor width (Table 4B), we found that highly docile sheep made more behavioural transitions in phase 3 than in phase 2 (Fig. 3B). Thus overall, wavelet analyses of longitudinal movements can be used as a proxy of sociability, and analyses of latitudinal movements give information about docility.

**Table 4:**
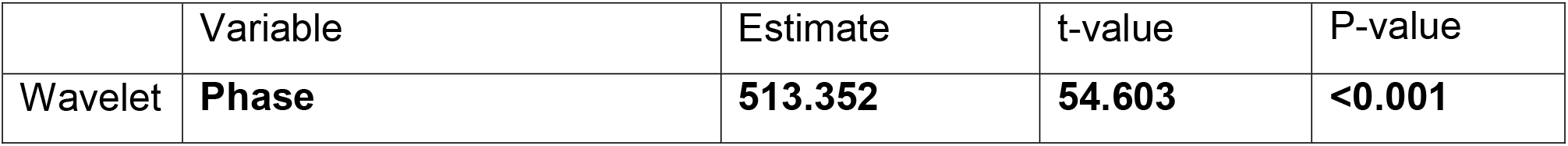
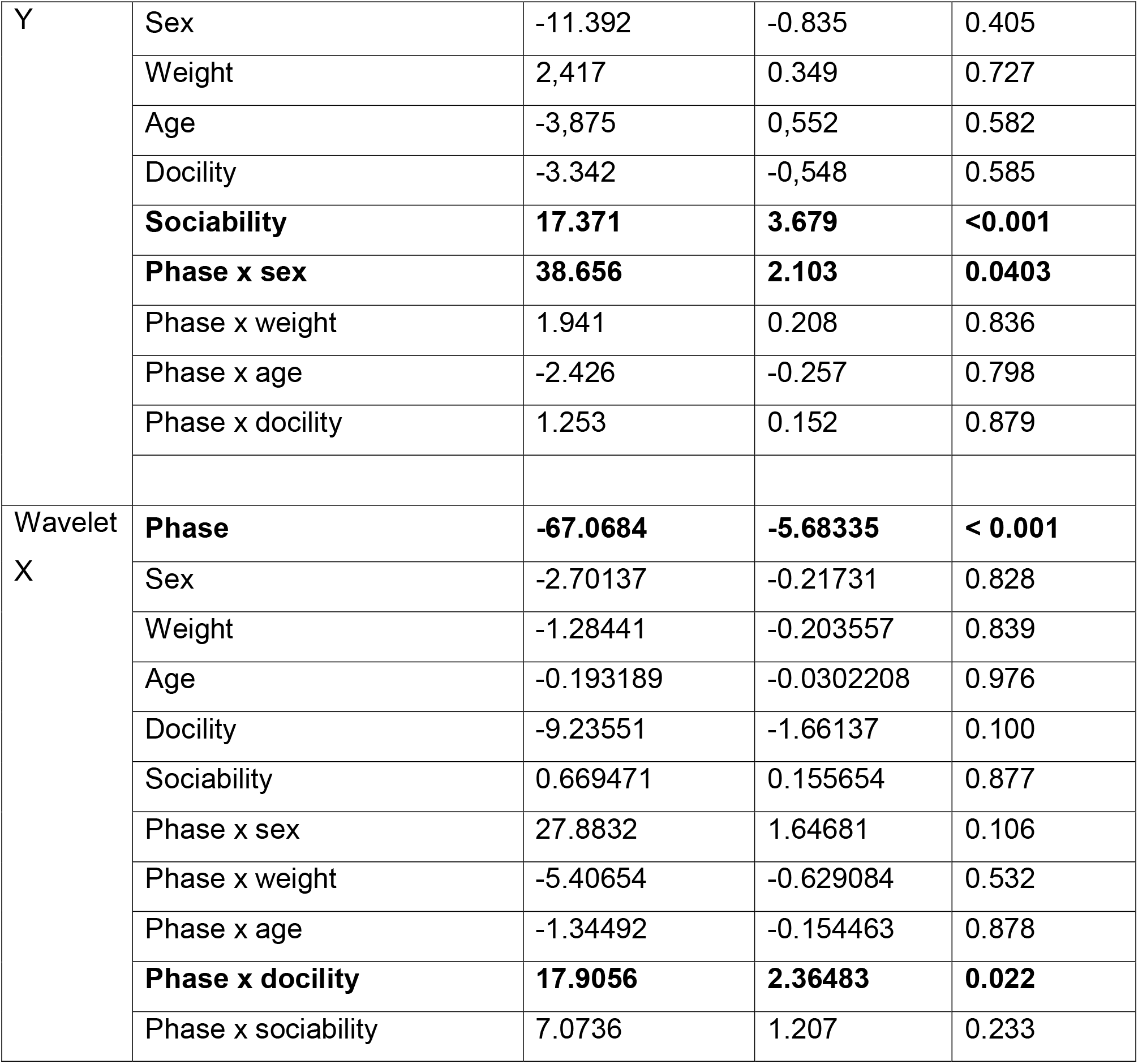
Wavelet analyses. Results of the best GLMM (Poisson family, after model selection – see details in Tables S2 and S3). The model tested the effects of phase, docility, weight, sex, age, sociability, and binary interactions of each variable with phase, on the number of wavelets. Lamb identity was included as a random factor. Significant effects (p<0.05) are shown in bold. Wavelet Y: longitudinal movements. Wavelet X: latitudinal movements.

#### Heatmap analyses: Detection of occupied space

We quantified spatial coverage by individual sheep (number of zones occupied in the corridor) using heatmaps. We tested the effects of the individual characteristics on the number of zones in which the sheep spent more than 200ms using GLMMs and model selection (Tables S4 and S5). When considering a grid with low spatial resolution, i.e. similar zone dimensions as with infrared cells (i.e., dimension: 0.6 × 1m; example Fig. 4A), we found that sheep used 2.37 times less space in phase 3 than in phase 2 (Table 5A). Larger individuals used more space than smaller ones (Table 5A). Increasing the spatial resolution of the grid, i.e. similar zone dimension as a lamb body size (i.e. dimension: 0.44 × 0.40 m; example Fig 4A), revealed that highly docile sheep used more space in phase 3 than in phase 2 (Table 5B). Young and highly social individuals used more space than old and poorly social individuals in both phases. Body size effect, however, was not significant anymore. Therefore, heatmap analyses are sensitive to grid size resolution. Decreasing grid size increased behavioural resolution.

**Table 5:** Heatmap analyses. Results of the best GLMM (Poisson family, after model selection – see details in Tables S4 and S5). The model tested the effects of phase, docility, weight, sex, age, sociability, and dual interactions of each variable with phase, on the number of areas where the lamb spent more than 1s. Sheep identity was included as a random factor. Significant effects (p<0.05) are shown in bold. A. Low spatial resolution. B. High spatial resolution.

|  | Variable | Estimate | t-value | P-value |
| --- | --- | --- | --- | --- |
| A | **Phase** | **-0.597701** | **-9.20941** | **< 0.001** |
|  | **Weight** | **0.0615106** | **1.99628** | **0.046** |
|  | Sociability | 0.0365622 | 1.63703 | 0.102 |
| B | Sex | 0.0188161 | 0.299427 | 0.765 |
|  | **Age** | **-0.0653912** | **-2.04381** | **0.041** |
|  | **Phase** | **-0.765708** | **-14.4685** | **< 0.001** |
|  | **Docility** | **-0.0837715** | **-2.665** | **0.008** |
|  | **Sociability** | **0.0480015** | **2.19218** | **0.029** |
|  | Weight | 0.0407016 | 1.27774 | 0.201 |
|  | **Phase x docility** | **0.0978634** | **2.15262** | **0.031** |

### Sheep behavioural phenotype

We explored whether the new behavioural measures extracted from the radar data could capture information from behavioural traits measured manually by the experimenter in the corridor test. We focused on docility and sociability.

We ran a PCA based on the eight behavioural measures extracted from the radar data in phase 2 and phase 3: proportion of fast movements (class 1) out of all movements (class 1 + class 2), longitudinal movements (wavelets Y), latitudinal movements (wavelets X), space coverage (heatmaps). We retained two PCs using the Kaiser-Guttman criterion (Kaiser, 1991). PC1 explained 36% of the variance and PC2 explained 22% of the variance. PC1 was positively associated with all behavioural variables (Fig. 5A). Sheep with high PC1 values moved more often faster, made more behavioural transitions, and used more zones than sheep with low PC1 values. We therefore interpreted PC1 as a “movement speed” variable. PC2 was positively associated with the four behavioural variables of phase 3 and negatively associated with the four behavioural variables of phase 2 (Fig. 5A). Sheep with high PC2 values showed a more important increase of time spent moving fast, of the frequency of behavioural transitions, and numbers of zones occupied from phase 2 to phase 3 than sheep with low PC2 levels. We interpreted PC2 as a variable of “movement increase between phases”.

**Figure 5:**
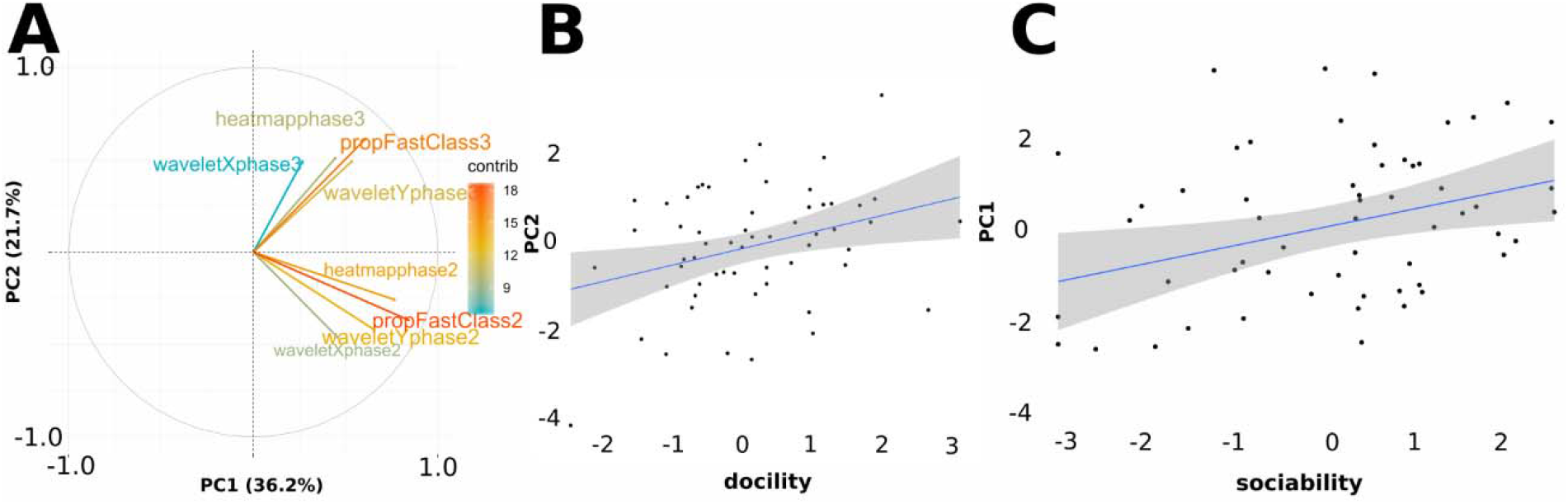
A. Correlations between the two first components (PCs) of the principal component analysis (PCA). Arrows represent the eight behavioural variables on PC1 (movement speed) and PC2 (movement increase between phases). Contribution of variables to the variance explained is colour coded. Each data point represents the PC1 and PC2 scores of a given lamb (N = 64). B. Relationship between PC1 and sociability. C. Relationship between PC2 and docility. Blue lines represent linear models (see main text). N = 64 sheep.

Sociability was significantly explained by PC1 (LM, PC1: estimate = 0.027, t = 2.552, p = 0.014; PC2: estimate = −0.104, t = −0.764, p = 0.448; Fig. 5B). Docility was significantly explained by PC2 (LM, PC1: estimate = −0.081, t = −0.943, p = 0.350; PC2: estimate = 0.283, t = 2.547, p = 0.014; Fig. 5C). Therefore, the automatically extracted radar data can capture inter-individual variability in sociability and docility traits usually measured by experimenters.

### Outdoor radar tracking

To demonstrate that our tracking system can be used in various experimental contexts, we tracked sheep in an outside corridor at a larger spatial scale (10 × 60m; Fig. 6A). We monitored the 2D trajectory of one sheep over a maximum distance of 45m (Fig. 6B). The presence of the man to induce sheep movement did not impair tracking (Fig. 6C).

**Figure 6.**
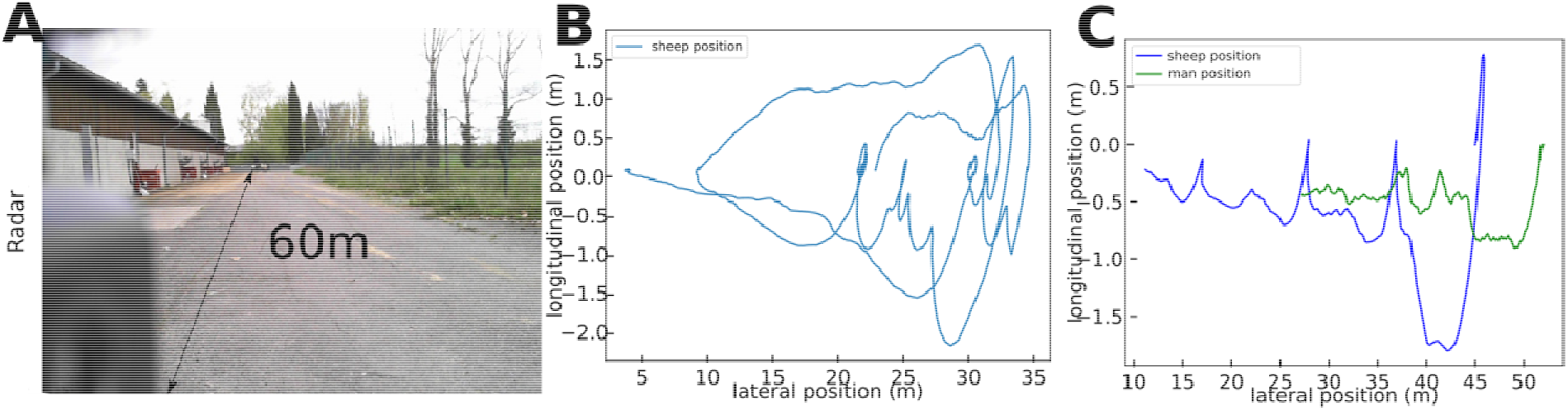
A. Picture of the outside corridor used for radar tracking of a sheep. The radar was positioned 60m from the end of the corridor. B. Example of trajectory of a sheep derived from radar data. C. Example of trajectory derived from radar data of a sheep (red) and a man (green) to induce the sheep movement.

## Discussion

Research in behaviour and ecology increasingly requires automated monitoring and annotation of animal movements for comparative quantitative analyses (Anderson and Perona, 2014; Brown and de Bivort, 2018). Here we introduced a radar tracking system suitable to study the 2D movements of sheep within a range of 45 m. The system is non-sensitive to light variations, compatible with real time data analyses, transportable, fast processing and adaptable to various species and experimental contexts. Moreover, it does not require to use tags or transponders for tracking the animals. It is therefore suitable for the collection of large sets of behavioural data in an automated way required in many areas of biological and ecological research.

FMCW radars were recently used to track sheep and pigs in 1D (Dore et al., 2020b; Henry et al., 2018) and bees in 3D (Dore et al., in press). Here, for the first time, we demonstrate the applicability of this approach to monitor 2D trajectories of untagged walking animals within a range of 45 meters with high spatial resolution. Radar acquisition system has several advantages over more conventional methods, and in particular video tracking, as it collects more data per seconds, requires less RAM, less processing time (e.g. does not require to train neural networks) and generates less false detection rates. It can therefore be applied for real-time detection of animal position. Importantly, the radar is not dependent on brightness and can be used for outside tracking over long distances by adjusting operating frequencies. It also enables the tracking of individualised animals without tags, based on the size and shape of the radar echoes of the different targets. Others methods can be used to estimate the sheep position. The main two methods are video detection (Bonneau et al., 2020), which can detect sheep in 2D up to 25m but with a precision of about 2m, and GPS detection (Gwyn et al., 1995), but this requires to animals.

Our application of radar-based tracking to behavioural phenotyping of sheep shows that the radar analysis is consistent with current manual or semi-automated analyses. we found that sheep tend to have a greater displacement in phase 2 than in phase 3 of the corridor test. This is consistent with previous work showing that sheep are more active when socially isolated, presumably as the result of them searching for social contact with conspecifics. These docile animals tend to move less when they are in contact with men. Importantly, the high resolution 2D trajectories obtained from the radar enabled to identify new behavioural estimators that could greatly benefit the fast and automated identification of behavioural phenotypes. These different estimators are not dependent on the radar tracking system per se, but requires to detect the position of the sheep with sufficient time resolution. For example, our application of unsupervised behavioural annotation to identify statistically significant behaviours by sheep in the corridor test showed that sheep exhibit less fast movements in phase 3 than in phase 2. Our utilisation of wavelet analyses revealed the occurrence of erratic displacements. The more the sheep was sociable the more it made erratic longitudinal movements. The more the sheep was docile, the more it made erratic latitudinal movements. We also analysed space occupation by sheep, showing that individuals exploit narrower areas in phase 3 than in phase 2. All these results are consistent with previous observations using manual or semi-automated recording methods. Most importantly, the combination of these new automatically computed estimators appears to be correlated with behavioural traits of interest that were until now measured manually by an experimenter during the corridor test or in complementary tests (e.g. carousel test). Therefore, in principle, our automated tracking and analysis system can be used for automated classification of animal behavioural profiles, which is a major issue for mass phenotyping in animal selection (Beausoleil et al., 2012). Note however that our pioneering study is based on relatively low sample sizes (64 individuals) and further measurements are needed to verify the biological trends observed on a much larger number of sheep.

Beyond genetic selection of farm animals, our system can be tuned to suit a large diversity of animal sizes and experimental contexts. Range and resolution of detection can be improved using different radars. For instance, we had to placed the radar at 1 meter from the corridor in order to track the entire corridor area. But with other antennas and a radar with larger field of view could have been placed at the edge of the corridor. Moreover, the detection was limited to a few meters but it is possible to detect a sheep at tens of meters using a radar operating at a lower frequency (24GHz) and/or transmitting higher electromagnetic power. It is also possible to improve radar detection by using more antennas. Indeed, by multiplying the number of antennas, we multiply the number of signal estimations and then the noise from the radar can be decreased. In the future, the same radar technology could be used to track individuals in groups over longer distances in open fields, for instance to explore the mechanisms underpinning social network structures and collective behaviour (Ginelli et al., 2015; King et al., 2012). Furthermore, it is possible to improve the processing of the radar signal for tracking large number of sheep simultaneously by using deep radar processing but this would require the use of a large amount of annotated data to train the neural networks (Huang et al., 2018). Individual tracking within groups could be improved with non-invasive passive tags that depolarize radar signal in specific directions (Lui and Shuley, 2006).

To conclude we demonstrate the feasibility of tracking a sheep in a restricted area using a FMCW radar. This detection is possible even if each wall of the corridor backscatters the transmitted electromagnetic signal. This radar can also be advantageously used to extract features that are relied to the movement of the sheep and can estimate if it is erratic, fast and the space occupied in the corridor. In contrast to other short-range methods, this 2D detection method does not require pre-annotated data and can be applied in real time. This flexibility holds considerable premises for tracking the behaviour of animals of various sizes and environments in a wide range of contexts and research fields.

## Acknowledgements

AD was funded by a PhD fellowship from the Council of the Région Occitanie (SIDIPAR Project). HA and ML also received support from grant of the French National Research Agency (ANR-DFG 3DNaviBee; ANR TERC BEE-MOVE).

## Supplementary materials

**Data S1:** Raw trajectories obtained from radar and video for each sheep.

**Table S1:**
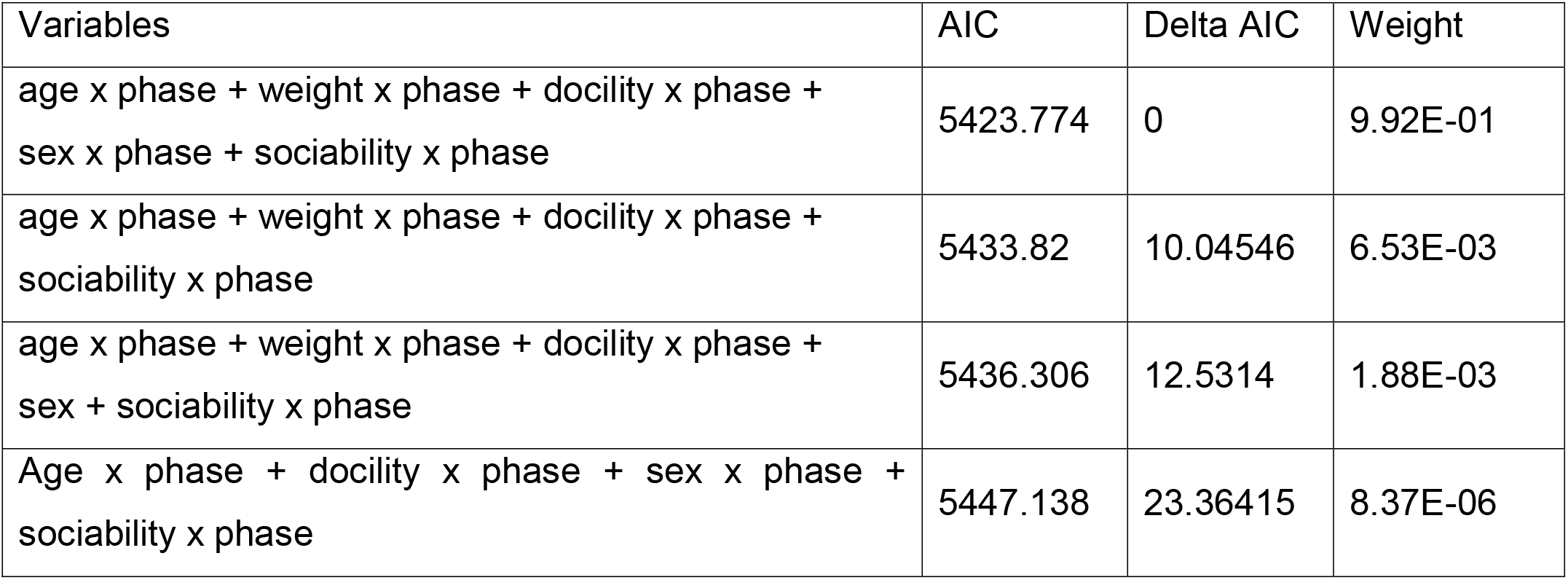
Model selection for behavioural class analyses. Null model, best model, second and third best models are displayed.

| Variables | AIC | Delta AIC | Weight |
| --- | --- | --- | --- |
| age x phase + weight x phase + docility x phase + sex x phase + sociability x phase | 5423.774 | 0 | 9.92E-01 |
| age x phase + weight x phase + docility x phase + sociability x phase | 5433.82 | 10.04546 | 6.53E-03 |
| age x phase + weight x phase + docility x phase + sex + sociability x phase | 5436.306 | 12.5314 | 1.88E-03 |
| Age x phase + docility x phase + sex x phase + sociability x phase | 5447.138 | 23.36415 | 8.37E-06 |

**Table S2:** Model selection for X wavelet analyses (latitudinal movements). Null model, best model, second and third best models are displayed.

| Variables | AIC | Delta AIC | Weight |
| --- | --- | --- | --- |
| age x phase + weight x phase + docility x phase + sex + sociability x phase | 1170.346 | 0 | 5.93E-01 |
| age + weight x phase + docility x phase + sex + sociability x phase | 1173.937 | 3.591115 | 9.85E-02 |
| age x phase + weight + docility x phase + sex + sociability x phase | 1174.289 | 3.942563 | 8.26E-02 |
| age x phase + weight + docility x phase + sex + sociability | 1174.586 | 4.239789 | 7.12E-02 |

**Table S3:**
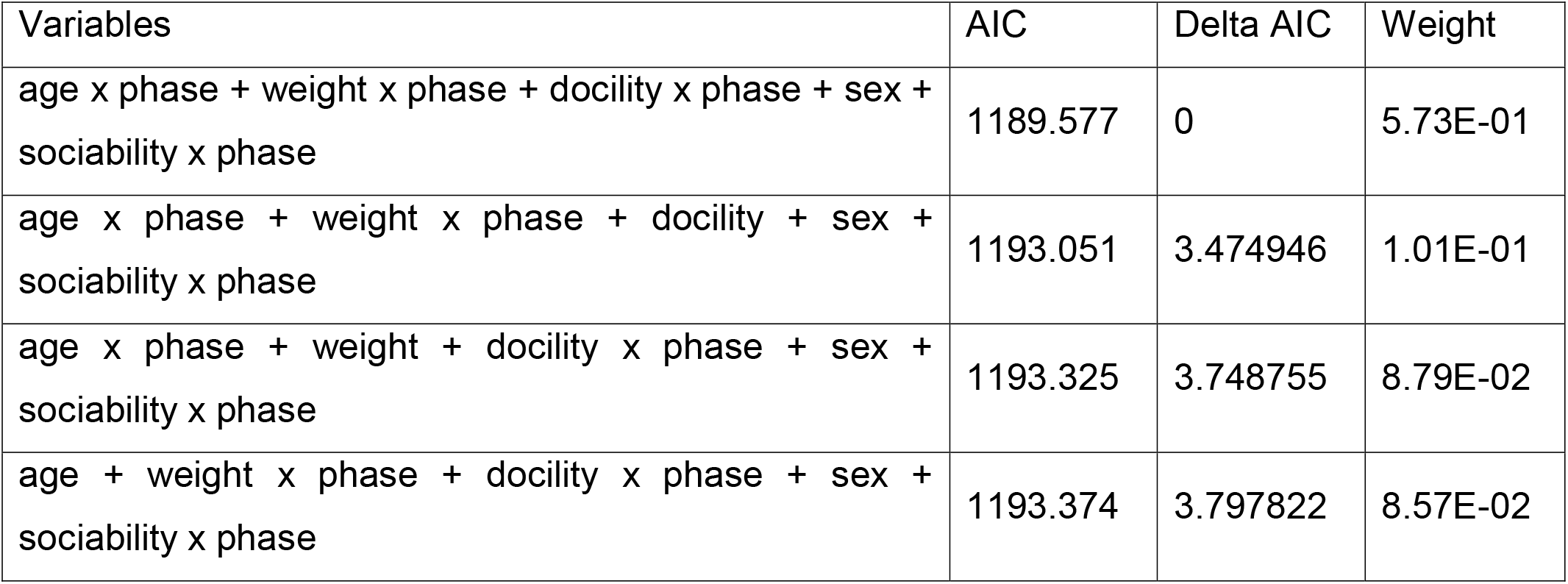
Model selection for Y wavelet analyses (longitudinal movements). Null model, best model, second and third best models are displayed.

**Table S4:** Model selection for heatmap analyses (low spatial resolution). Null model, best model, second and third best models are displayed.

| Variables | AIC | Delta AIC | Weight |
| --- | --- | --- | --- |
| weight x phase + sociability x phase | 544.1552 | 0 | 1.35E-01 |
| weight x phase + sex x phase + sociability x phase | 546.3065 | 2.151297 | 4.61E-02 |
| sex x phase + sociability x phase | 546.376 | 2.220815 | 4.46E-02 |
| weight x phase + sociability x phase + docility x phase | 546.569 | 2.413782 | 4.05E-02 |

**Table S5:** Model selection for heatmap analyses (high spatial resolution). Null model, best model, second and third best models are displayed.

| Variables | AIC | Delta AIC | Weight |
| --- | --- | --- | --- |
| age + docility x phase + weight x phase + sociability | 664.2827 | 0 | 0.058719217 |
| age + docility x phase + weight + sociability | 664.9303 | 0.6476525 | 0.042476067 |
| age + docility x phase + sociability x phase | 665.5479 | 1.2652317 | 0.031191675 |
| docility x phase + weight x phase + sociability x phase | 665.8718 | 1.5891709 | 0.026527492 |

## References

Anderson, D.J., Perona, P., 2014. Towards a science of computational ethology. Neuron 84, 18–31.

Balanis, C.A., 2011. Modern antenna handbook, John Wiley&Sons. ed.

Bates, D., Maechler, M., Bolker, B., Waleker, S., 2014. lme4: linear mixed-effects models using Eigen and S4. R package version 1.

Beausoleil, N.J., Blache, D., Stafford, K.J., Mellor, D.J., Noble, A.D.L., 2012. Selection for temperament in sheep: Domain-general and context-specific traits. Applied Animal Behaviour Science 139.

Boissy, A., Bouix, J., Orgeur, P., Poindron, P., Bibe, B., Le Neindre, P., 2005. Genetic analysis of emotional reactivity in sheep: effects of the genotypes of the lambs and of their dams. Genetics Selection Evolution 37, 381–401.

Bonneau, M., Vayssade, J.A., Troupe, W., Arquet, R., 2020. Outdoor animal tracking combining neural network and time-lapse cameras. Computers and Electronics in Agriculture 168.

Branson, K., Robie, A.A., Bender, J., Perona, P., Dickinson, M.H., 2009. High-throughput ethomics in large groups of Drosophila. Nat Methods 6, 451–457.

Brown, A.E.X., de Bivort, B., 2018. Ethology as a physical science. Nat Phys 14, 653–657.

Burnham, K.P., Anderson, D.R., 2002. Model selection and multimodel inference: a practical information-theoretic approach, 2nd Edition. ed. Springer, New York, NY.

Cadahia, L., López-López, P., Urios, V., Negro, J.J., 2010. Satellite telemetry reveals individual variation in juvenile Bonelli’s eagle dispersal areas. European Journal of Wildlife Research 56.

Canario, L., Mignon-Grasteau, S., Dupont-Nivet, M., Phocas, F., 2013. genetics of behavioural adaptation of livestock to farming conditions. Animal 7, 357–377.

Dell, A.I., Bender, J.A., Branson, K., Couzin, I.D., de Polavieja, G.G., Noldus, L., Pérez-Escudero, A., Perona, P., Straw, A.D., Wikelski, M., Brose, U., 2014. Automated image-based tracking and its application in ecology. Trends Ecol Evol 29, 417–428. https://doi.org/10.1016/j.tree.2014.05.004

Dietlein, C.R., Bjarnason, J.E., Grossman, E.N., Popovic, Z., 2008. Absorption, transmission, and scattering of expanded polystyrene at millimeter-wave and terahertz frequencies. Passive Millimeter-Wave Imaging Technology XI 6948–69480.

Dore, A., Henry, D., Aubert, H., Lihoreau, M., 2020a. 3D trajectories of multiple untagged flying insects from millimetre-wave beamscanning radar. Presented at the IEEE International Symposium on Antennas and Propagation and Notth American Radio Science Meeting, Québec, Canada.

Dore, A., Henry, D., Aubert, H., Lihoreau, M., in press. 3D trajectories of multiple untagged flying insects from millimetre-wave beamscanning radar. IEEE Antennas and Propagation.

Dore, A., Lihoreau, M., Billon, Y., Ravon, L., Reignier, S., Bailly, J., Bompa, J.F., Ricard, E., Aubert, H., Henry, D., Canario, L., 2020b. Millimeter-wave Radars for the Automatic Recording of Sow Postural Activity. Presented at the 71st Annual Meeting of European Federation of Animal Science, Porto, Portugal.

Everingham, M., Van Gool, L., Williams, C.K.I., Winn, J., Zisserman, A., 2012. Results 2012. The PASCAL Visual Object Classes Challenge 2012 (VOC2012).

Ginelli, F., Peruani, F., Pillot, M.H., Chaté, H., Theraulaz, G., Bon, R., 2015. Intermittent collective dynamics emerge from conflicting imperatives in sheep herds. Proc Natl Acad Sci USA 112, 12729–12734.

Gwyn, R., Williams, A., Last, J.D., Penning, P.D., Mark Rutter, S., 1995. A Low-Power Postprocessed DGPS System for Logging the Locations of Sheep on Hill Pastures. Navigation 42.

Haderer, A., Wagner, C., Feger, R., Stelzer, A., 2008. A 77-GHz FMCW front-end with FPGA and DSP support, in: 2008 International Radar Symposium. pp. 1–6.

Hazard, D., Moreno, C., Foulquié, D., Delval, D., François, D., Bouix, J., Sallé, G., Boissy, A., 2014. Identification of QTLs for behavioral reactivity to social separation and humans in sheep using the OvineSNP50 BeadChip. BMC Genomics 15, Artilce 778.

Hazard, D., Bouix, J., Chassier, M., Delval, E., Foulquié, D., Fassier, T., Bourdillon, Y., François, D., Boissy, A., 2016. Genotype by environment interactions for behavioral reactivity in sheep. J Anim Sci 94, 1459–1471.

Henry, D., Aubert, H., Ricard, E., Hazard, D., Lihoreau, M., 2018. Automated monitoring of livestock behavior using frequency-modulated contnuous-wave radars. Prog Electromagn Res 69, 151–160.

Huang, H., Gui, G., Sari, H., Adachi, F., 2018. Deep learning for super-resolution DOA estimation in massive MIMO systems. Presented at the IEEE 88th Vehicular Technology Conference (VTC-Fall).

Idris, A., Moors, E., Budnick, C., Herrmann, A., Erhardt, G., Gauly, M., 2011. Is the establishment rate and fecundity of Haemonchus contortus related to body or abomasal measurements in sheep? Animal 5, 1276–1282.

Kaiser, H.F., 1991. Coefficient alpha for a principal component and the Kaiser-Guttman rule. Psychol Rep 68, 855–858.

King, A.J., Wilson, A.M., Wilshin, S.D., Lowe, J., Haddadi, H., Hailes, S., Morton, J., 2012. Selfish-herd behaviour of sheep under threat. Curr Biol 22, R561–562.

Ligout, S., Foulquié, D., Sèbe, F., Bouix, T., Boissy, A., 2011. Assessement of sociality in farm animals: the use of the arena test in lambs. Appl Anim Behav Sci 135, 57–62.

Lui, H.S., Shuley, N., 2006. Resonance based radar target identification with multiple polarizations. Presented at the EEE Antennas and Propagation Society International Symposium.

Morand-Ferron, J., Cole, E.F., Quinn, J.L., 2015. Studying the evolutionary ecology of cognition in the wild: a review of practical and conceptual challenges. Biol Rev 91, 367–389.

Pérez-Escudero, A., Vicente-Page, J., Hinz, R.C., Arganda, S., de Polavieja, G.G., 2014. idTracker: tracking individuals in a group by automatic identification of unmarked animals. Nat Methods 11, 743–748. https://doi.org/10.1038/nmeth.2994

Phocas, F., Boivin, X., Sapa, J., Trillat, G., Boissy, A., Le Neindre, P., 2006. Genetic correlations between temperament and breedng traits in Limousin heifers. Animal Science 82, 805–811.

Poirier, J.R., Aubert, H., Jaggard, D.L., 2009. Lacunarity of rough surfaces from the wavelet analysis of scattering data. IEEE Transactions on Antennas and Propagation 57, 2130–2136.

R Core Team, 2014. R: a Language and Environment for Statistical Computing, R Foundation for Statistical Computing. ed. Vienna, Austria.

Redmon, J., Divvala, S., Girshick, R., Farhadi, A., 2016. You Only Look Once: Unified, Real-Time Object Detection. Proceedings of the IEEE Conference on Computer Vision and Pattern Recognition 779–788.

Reynolds, D.A., Rose, R.C., n.d. Robust text-independent speaker identification using Gaussian mixture speaker models. IEEE Transactions on Speech and Audio Processing 3, 72–83.

Riley, J.R., Smith, A.D., Reynolds, D.R., Edwards, A.S., Osborne, J.L., Williams, I.H., Carreck, N.L., Poppy, G.M., 1996. Tracking bees with harmonic radar. Nature 379, 29–30. https://doi.org/10.1038/379029b0

Romero-Ferrero, F., Bergomi, M.G., Hinz, R.C., Heras, F.J.H., de Polavieja, G.G., 2019. idtracker.ai: tracking all individuals in small or large collectives of unmarked animals. Nature Methods 16, 179–182.

Ryan, H., 1994. Ricker, Ormsby, Klauder, Butterworth - A choice of wavelets. CSEG Recorder 8–9.

Simon, W., Klein, T., Litschke, O., 2014. Small and light 24 GHz multi-channel radar. Presented at the IEEEAntennas and Propagation Society International Symposium (APSURSI).

Tomkiewicz, S.M., Fuller, M.R., Kie, J.G., Bates, K.K., 2010. Global positioning system and associated technologies in animal behaviour and ecological research. Philosophical Transactions of the Royal Society B. https://doi.org/10.1098/rstb.2010.0090

Voulodimos, A.S., Patrikakis, C.Z., Sideridis, A.B., Ntafis, V.A., Xylouri, E.M., 2010. A complete farm management system based on animal identification using RFID technology. Computers and Electronics in Agriculture 70, 380–388. https://doi.org/10.1016/j.compag.2009.07.009

